# High-throughput screen using fluorescence lifetime detects compounds that modulate myosin-binding protein C interactions with actin

**DOI:** 10.1101/2021.03.24.436879

**Authors:** Thomas A. Bunch, Piyali Guhathakurta, Victoria C. Lepak, Andrew R. Thompson, Rhye-Samuel Kanassatega, Anna Wilson, David D. Thomas, Brett A. Colson

## Abstract

Cardiac myosin-binding protein C (cMyBP-C) interacts with actin and myosin to modulate cardiac contractility. These interactions are regulated by cMyBP-C phosphorylation. Heart failure patients often have decreased cMyBP-C phosphorylation and phosphorylation in model systems appears to be cardioprotective for heart failure. Therefore, cMyBP-C is a potential target for heart failure drugs that mimic phosphorylation and/or perturb its interactions with actin/myosin.

We have used a novel fluorescence lifetime-based assay to identify small-molecule inhibitors of actin-cMyBP-C binding. Actin was labeled with a fluorescent dye (Alexa Fluor 568, AF568) near its cMyBP-C binding sites. When combined with cMyBP-C N-terminal fragment, C0-C2, the fluorescence lifetime of AF568-actin decreases. Using this reduction in lifetime as a readout of actin binding, a high-throughput screen of a 1280-compound library identified 3 reproducible Hit compounds that reduced C0-C2 binding to actin in the micromolar range. Binding of phosphorylated C0-C2 was also blocked by these compounds. That they specifically block binding was confirmed by a novel actin-C0-C2 time-resolved FRET (TR-FRET) binding assay. Isothermal titration calorimetry (ITC) and transient phosphorescence anisotropy (TPA) confirmed that the Hit compounds bind to cMyBP-C but not to actin. TPA results were also consistent with these compounds inhibiting C0-C2 binding to actin. We conclude that the actin-cMyBP-C lifetime assay permits detection of pharmacologically active compounds that affect cMyBP-C’s actin binding function. TPA, TR-FRET, and ITC can then be used to understand the mechanism by which the compounds alter cMyBP-C interactions with actin.

---

Cardiac myosin-binding protein C (cMyBP-C, **Fig. 1**) has been identified as a therapeutic target for systolic or diastolic dysfunction in heart failure and cardiomyopathy. cMyBP-C may contribute to heart failure due to hypertrophic cardiomyopathy (HCM) or dilated cardiomyopathy (DCM).

**Figure 1.**
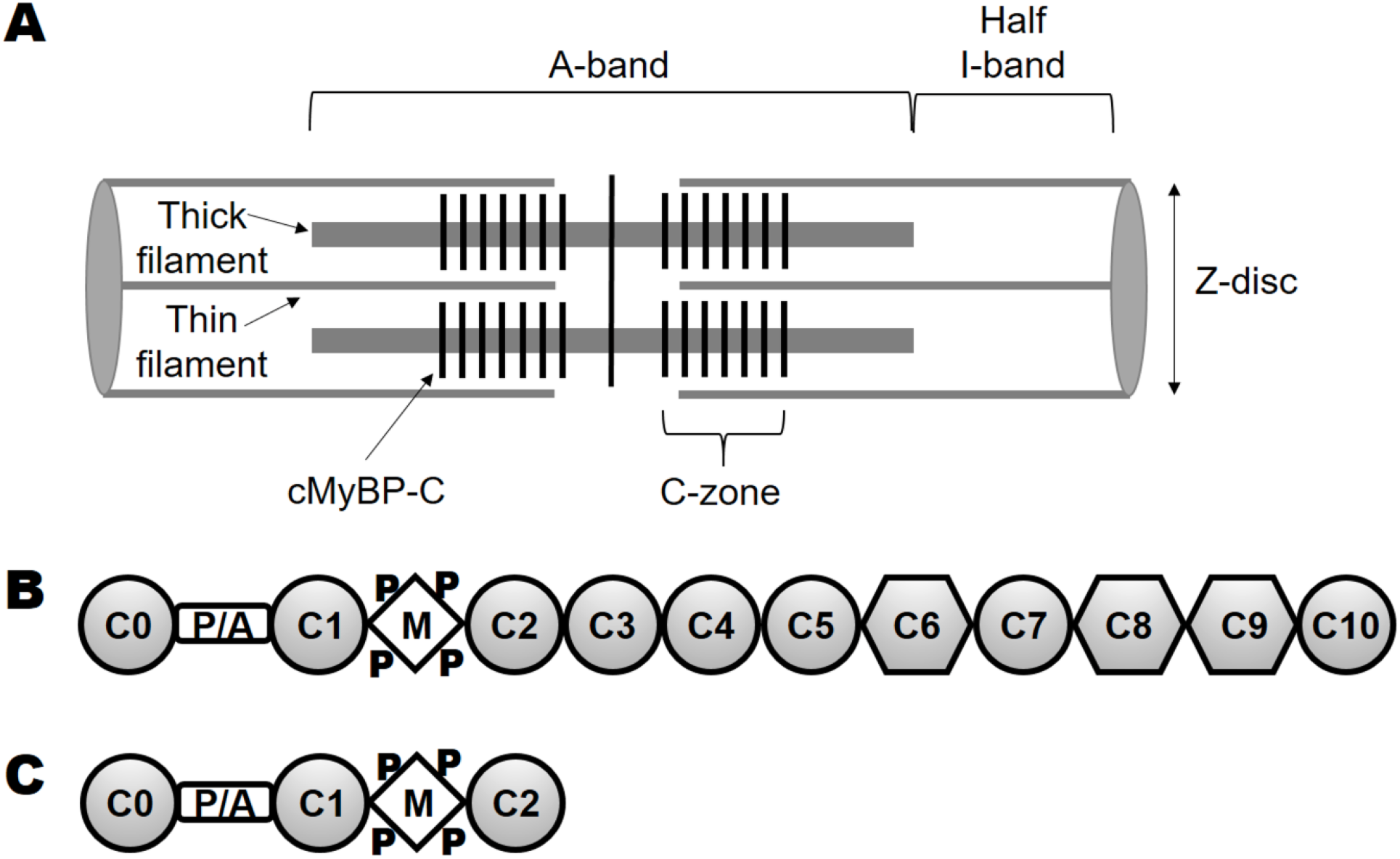
cMyBP-C organization. ***(A)*** The sarcomere spans from Z-disc to Z-disc with the A-band containing thick filaments and the I-band containing actin filaments. Force is generated by myosin and actin in the thin/thick filament overlap portion of the A-band. cMyBP-C molecules (black vertical stripes) are present in the C-zones toward the center of the A-band. The C-zones overlap with thin filaments (except at very long sarcomere lengths). ***(B)*** Full-length cMyBP-C domains C0 through C10. Ig-like domains are shown as circles and fibronectin type-III domains are shown as hexagons. ***(C)*** N-terminal domains C0 through C2 (C0-C2), containing the proline alanine rich linker (P/A) and the M-domain (M) that contains phosphorylation sites (P).

Hypertrophic cardiomyopathy (HCM) is a common heart disease, affecting around 1:500 individuals, for which there are no therapeutic treatments (1). In HCM the heart typically becomes enlarged, hypercontractile, and unable to relax effectively. The systolic contraction is preserved, but diastolic relaxation is diminished. HCM is associated with heart failure, arrhythmias, and cardiac death at any age. The primary cause of HCM is most often a missense mutation in one of several sarcomeric proteins, or the truncation of cMyBP-C. In addition to cMyBP-C, the sarcomeric proteins associated with HCM include those associated with the thick (myosin) and thin (actin) filaments, including myosin, troponin, tropomyosin, leiomodin, and titin.

Dilated cardiomyopathy (DCM) results from missense mutations in some of the same genes as for HCM (but not cMyBP-C truncations). It leads to dilation of the ventricular chambers and defects in systolic contraction.

Numerous studies have demonstrated that increasing or decreasing PKA-mediated phosphorylation of cMyBP-C allows for tuning cardiac contraction and relaxation through phosphorylation-sensitive interactions with actin and myosin. Recent work by us (2) and others (3) suggests that phosphorylation causes a rearrangement of N-terminal cMyBP-C structure that reduces binding to actin and myosin to modulate contraction and relaxation. Decreases in phosphorylation of cMyBP-C have been observed in heart failure patients, including those affected by mutations in sarcomeric proteins other than cMyBP-C. Therefore, targeting cMyBP-C with drugs that mimic phosphorylation and that modulate its binding to actin or myosin is a promising approach to improve cardiac muscle function in heart failure and cardiomyopathy. Recently, cardiac-specific, small-molecule effectors of myosin have been developed to activate or inhibit cardiac muscle contraction in heart failure and hypertrophic cardiomyopathy (4,5). While some ongoing human clinical trials are promising, alternative strategies, such as targeting cardiac specific actomyosin accessory proteins (i.e. cMyBP-C) may prove valuable as well. Despite strong evidence implicating cMyBP-C as an ideal target for therapy, no cMyBP-C small-molecule modulator has been identified. A critical barrier to progress in this area is that high-throughput screening (HTS) assays based on cMyBP-C functional activities have not been available.

Current *in vitro* actin or myosin binding assays, such as cosedimentation, are labor intensive and low-throughput, limited in the number of samples that can be tested. Recent development of fluorescence lifetime plate readers (6) now allows for high-throughput (~3 minutes/microplate), high-precision (picoseconds for lifetime and nanometers for distances) assays that are scalable (in-plate, 1,536-well plates, detection) and reduce labor.

We recently developed a fluorescence lifetime HTS assay for detecting the interaction of F-actin with N-terminal cMyBP-C domains C0-C2 (7). The time-resolved fluorescence (TR-F) assay reported a significant reduction in lifetime of the dye attached to 1 μM actin at Cys-374 when incubated with C0-C2. This reduction was concentration-dependent and correlated with actin binding of C0-C2 as measured by cosedimentation assays. The high precision of the TR-F assay was demonstrated by resolving significant changes in binding due to phosphorylation and mutations relevant to HCM and function.

In the present study, we have used this TR-F assay, using a fluorescence lifetime plate reader (see Experimental Procedures) to screen the Library of Pharmacologically Active Compounds (LOPAC), which is a diverse collection of 1280 pharmacologically active compounds that includes all major drug target classes, inhibitors, pharma-developed tools, and approved drugs. To reduce interference with compounds exhibiting fluorescence at lower wavelengths (8,9), we labeled actin with the red-shifted fluorescent dye, Alexa Fluor 568 (AF568), which was excited by a 532-nm microchip laser. This screen reproducibly identified 3 compounds that appeared to reduce MyBP-C binding to actin in the micromolar range. We then tested these “Hit” compounds for dose-dependence in the TR-F assay and further evaluated their efficacy in biochemical and biophysical secondary assays. To verify compound-target engagement, isothermal titration calorimetry (ITC) determined the micromolar affinity of the three Hit compounds to cMyBP-C. Transient phosphorescence anisotropy (TPA) provided further proof that the Hit compounds did not bind to actin and that they inhibited C0-C2-actin interactions. Finally, we developed a novel TR-FRET assay as an alternative method to monitor C0-C2 binding to actin. We used this TR-FRET assay to confirm the effects of the identified Hits on actin binding.

This is this first HTS assay to identify compounds that specifically bind to human cMyBP-C and modulate its interactions with actin. The results of this study provide proof-of-concept validation for the TR-F assay in identifying effective cMyBP-C-specific modulators of actomyosin function. This new tool will be valuable in the development of new therapies for cardiac muscle diseases.

## Results

### Actin-cMyBP-C TR-F biosensor

We have previously described a TR-F based C0-C2-actin binding assay utilizing IAEDANS-labeled actin. The lifetime, τ, of IAEDANS attached to actin was reduced upon C0-C2 binding. This provided a rapid way to monitor binding in a multi-well plate format conducive to screening libraries of compounds that modulate actin-cMyBP-C binding (7). Unfortunately, a high percentage of compounds fluoresce at the shorter wavelength used to excite IAEDANS (8). To decrease this problem, we tested the TR-F response of C0-C2 binding to actin, where the actin was labeled with probes that are excited at longer wavelengths, permitting the use of a longer-wavelength laser (8).

Time-resolved fluorescence (TR-F) decays of Alexa Fluor 546 (AF546) and Alexa Fluor 568 (AF568)-labeled actin in the presence of cMyBP-C N-terminal fragment (C0-C2) (**Fig. 2A**) were analyzed by one-exponential fitting (see *Experimental Procedures*) to determine fluorescence lifetime (**Fig. 2B-C** and **Fig. S1A** in *Supplemental Figures*). Both probes on actin showed lifetime decreased with increasing [C0-C2], indicating binding. This effect was reduced with phosphorylated C0-C2, indicating less binding (**Fig. 2D**). As AF568 showed a greater change in lifetime upon C0-C2 binding to F-actin, we used this probe for the HTS screen. Analysis of TR-F binding curves with 1 μM AF568-actin revealed that at 2 μM C0-C2 the lifetime is highly sensitive to phosphorylation by PKA (**Fig. 2D**). Since our goal was to detect compounds that show differences in binding similar to those detected upon PKA phosphorylation, we selected these concentrations of AF568-actin (1 μM) and C0-C2 (2 μM) to perform the screen. From cosedimentation assays (7), we know that these correspond to binding ratios of ~1:7 C0-C2:actin, which is similar to that found in the muscle sarcomere.

**Figure 2.**
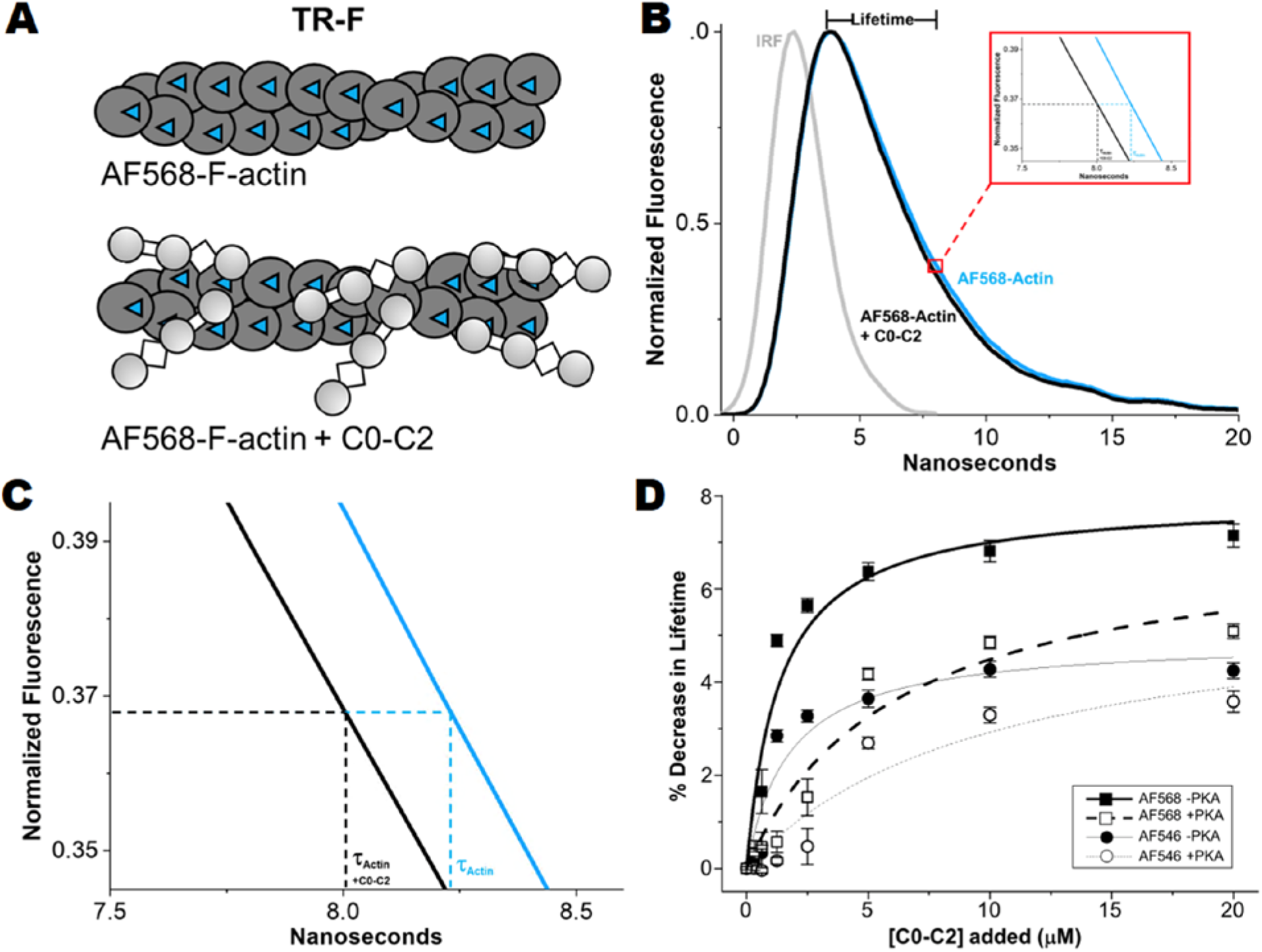
TR-F of AF568-F-actin is reduced upon C0-C2 binding. ***(A)*** Cartoon rendering of F-actin with AF568 (blue triangles) attached to each monomer (AF568-actin, dark grey circles) and the same upon binding of C0-C2. ***(B)*** Fluorescence waveforms of AF568-actin (blue line) and the same in the presence of 20 μM C0-C2 (black line) normalized to maximal fluorescence intensity (waveforms without normalization can be found in **Fig. S1A** in *Supplemental Figures*). Lifetime (τ) is the time at which the peak fluorescence decays to ~0.37 (1/e). Red box shows expansion of the x-axis near the lifetimes of the AF568- actin. ***(C)*** Further expansion of the red box area in B. ***(D)*** Comparison of AF546- (circles, thin grey lines) and AF568-actin (squares, thick black lines) binding curves for 0-20 μM C0-C2 without (solid circles/squares, solid lines) and with PKA phosphorylation (open circles/squares, dashed lines).

### High-throughput screening of LOPAC library to identify compounds that modulate actin-cMyBP-C TR-F

We performed HTS using LOPAC. The screen was performed in duplicate with two different preparations of AF568-actin and C0-C2 samples. Each screen tested the effect of the compounds on the lifetime of AF568-actin and the effect of the compounds on the lifetime of AF568-actin in the presence of C0-C2 (AF568-actin:C0-C2). The *Z’* factor for this screen was calculated as 0.5 ± 0.1 using DMSO-only controls, validating the robustness of this HTS assay (see Experimental Procedures). Compounds that altered the lifetime of AF568-actin alone by more than 4 standard deviations (SD) of the mean of controls (no compound) were excluded from the analysis of the AF568-actin:C0-C2 results. This removed 210 and 106 compounds from the first and second screen, respectively. The change in lifetime of AF568-actin due to effects of each compound on AF568-actin:C0-C2 complex was determined by subtracting the effects of each compound on 568-actin alone from the effects on AF568-actin:C0-C2. Since C0-C2 binding reduced AF568-actin lifetime, compounds interfering with C0-C2 binding to actin increased the lifetime of AF568-actin:C0-C2. Positive changes from both screens were sorted, ranked and compared. The top 25 compounds revealed 3 compounds (designated “Hits”) in common and 22 that were not repeatable (**Tables S1** and **S2** in *Supplemental Tables*). Two additional sub-optimal (explained in *Experimental Procedures*) screens of the same library also yielded only these 3 Hit compounds in the top 25 preliminary hits. Each of these had 22 false positives that were not identified in any of the other screens (**Tables S3** and **S4** in *Supplemental Tables*). Thus, the Hit rate was 0.23%.

Suramin showed the greatest effect, with an average change in lifetime that was 6.5 SD above the median (of all compounds not excluded due to large effects on AF568-actin alone); NF023 (a suramin analog) was 3.8 SD above the median; and aurintricarboxylic acid (ATA) was 3.0 SD above the median. These 3 Hit compounds were tested further in orthogonal assays.

### TR-F dose-response assay

Using the same conditions as in the primary screen, we determined the concentration-dependence of each of the 3 Hit compounds on the AF568-actin lifetime effects. As observed in the screen, all 3 Hit compounds had little effect on AF568-actin alone (**Fig. 3A**). C0-C2 reduced the lifetime of AF568, and all 3 Hit compounds, in a concentration-dependent manner, returned AF568 to the lifetime observed for AF568-actin alone (**Fig. 3A**). This is consistent with all 3 Hit compounds inhibiting C0-C2 binding to actin. The apparent EC_50_ (concentration of compound needed for half-maximal effect) of the Hit compounds as measured by TR-F is 5-20 μM (**Fig. 3, Table 1**). In a separate dose-response assay utilizing 3 μM C0-C2, we compared the effects of the compounds on nonphosphorylated and phosphorylatated C0-C2. When AF568-actin was bound to C0-C2 phosphorylated by PKA, a decreased lifetime of AF568 was observed (though the decrease was smaller than with nonphosphorylated C0-C2) and all 3 Hit compounds again returned AF568 lifetime to that of AF568-actin alone (**Fig. 3B**).

**Figure 3.**
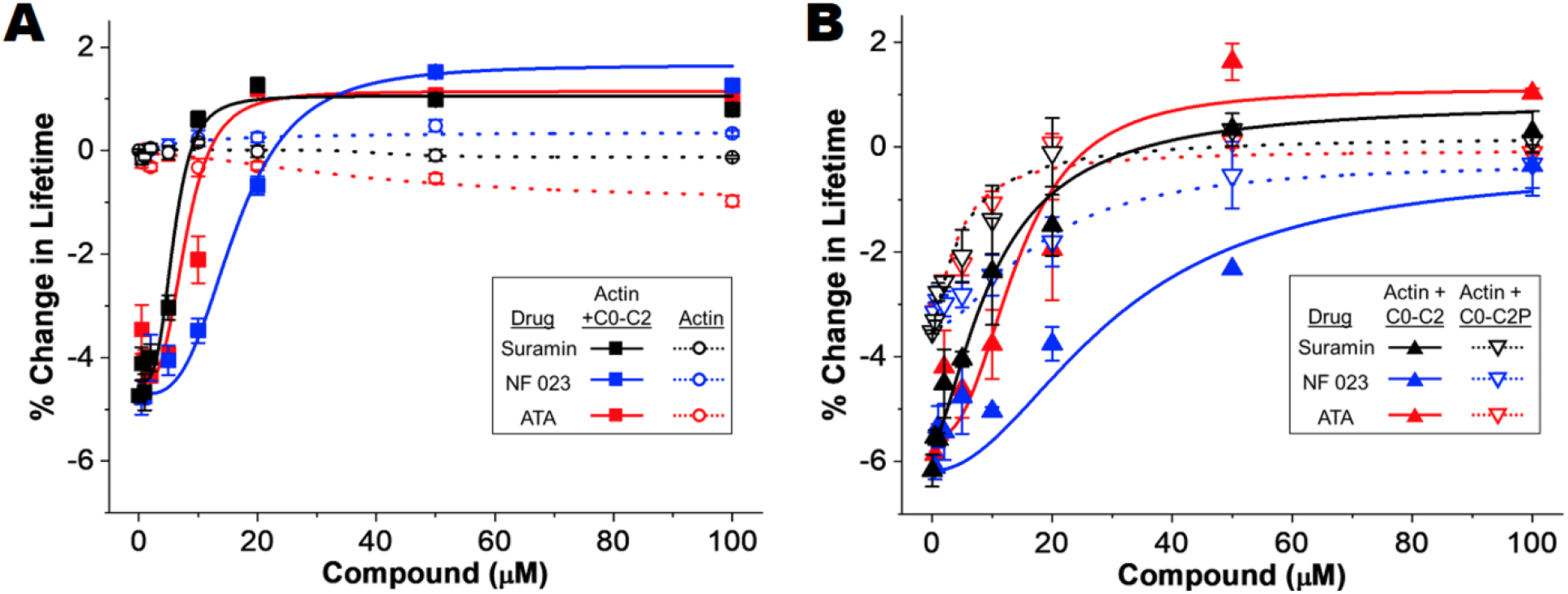
Concentration-response curves for Hit compounds on AF568-actin-C0-C2 TR-F. Percent change in AF568 Lifetime (relative to AF568-actin alone) for ***(A)*** 1 μM AF568-actin (open circles, dotted lines) and 1 μM AF568-actin with 2 μM C0-C2 (solid squares, solid lines) incubated with increasing concentrations of compounds (0-100 μM) (n=4 for each point) ***(B)*** AF568-actin (1 μM):C0-C2 (3 μM, closed triangles, solid lines) or AF568-actin (1 μM):phosphorylated (by PKA) C0-C2 (C0-C2P, 3 μM, open triangles, dotted lines), incubated with increasing amounts of Hit compounds (0-100 μM) (n=2 from separate actin and C0-C2 preparations). Average and SE for each data point are shown.

**Table 1:** Summary of Hit compound effects on the dose-response of AF568-actin-C0-C2 TR-F (left) and ITC (right) parameters of compound binding to C0-C2.

| Compound | TR-F | TR-F | ITC | ITC | ITC | ITC | ITC |
| --- | --- | --- | --- | --- | --- | --- | --- |
|  | EC<sub>50</sub> (μM) | n | K<sub>d</sub> (μM) | n | ΔH | ΔG | ΔS |
| Suramin | 5.9 ± 1.2 | 3.6 | 16.6 ± 1.0 | 3.4 | -17.4 | -26.8 | 32.3 |
| NF023 | 16.3 ± 1.2 | 3.2 | 42.0 ± 3.4 | 2.1 | -30.2 | -24.6 | -19.2 |
| ATA | 7.9 ± 1.9 | 3.4 | 25.9 ± 2.2 | 3.1 | -10.2 | -25.8 | 52.9 |

Average data are provided as mean ±SE and each experiment was done with 2 separate protein preparations.

### ITC: Compounds bind to C0-C2

Potential binding of these 3 Hit compounds to cMyBP-C was evaluated by Isothermal Titration Calorimetry (ITC). ITC titration curves of C0-C2 with each of the 3 identified compounds are shown in **Fig. 4A-C**. Very clear binding curves with saturation at an excess of C0-C2 indicates that there is direct interaction between the Hit compounds and C0-C2. Fitting the binding curves in **Fig. 4A-C** (and replicates done with a separate preparation of C0-C2) yielded dissociation constants of 15-75 μm and stoichiometries of C0-C2:compound of 1:2-3 (**Table 1**). For suramin, but not NF023 or ATA, weak interactions with actin alone (K_d_=~200 μM) were noted (**Fig. S2** in *Supplemental Figures*).

**Figure 4.**
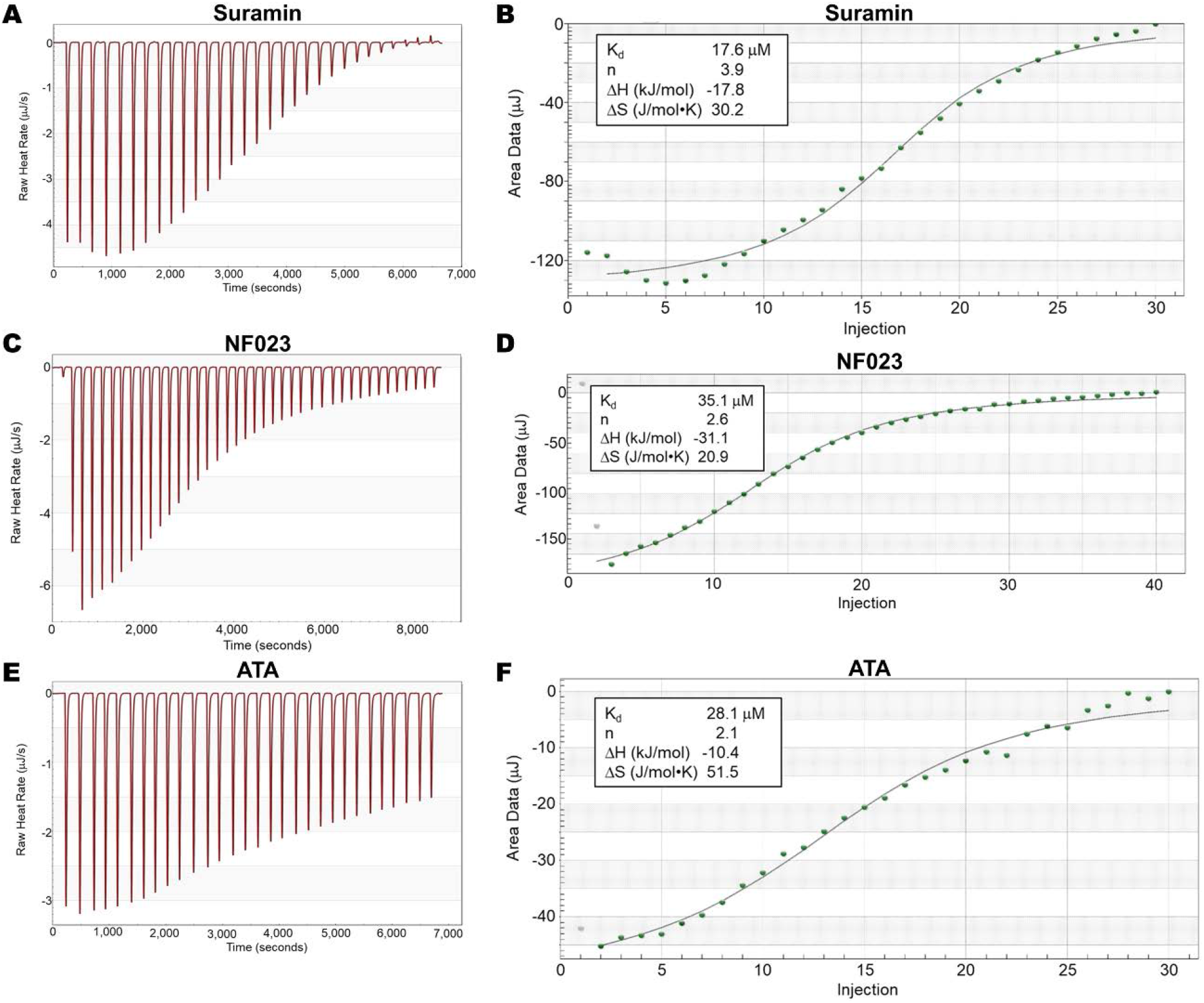
Isothermal Titration Calorimetry (ITC) of Hit compounds binding to C0-C2. ***(A)*** Titration of 100 μM C0-C2 (in 350 μl) with 30 2.5 μl injections of 3,000 μM Suramin. ***(B)*** Fitted ITC binding curve for Suramin binding. ***(C and D)*** The same for 2,500 μM NF023 (40 injections). ***(E and F)*** the same for 2,000 μM ATA (30 injections). The binding of each Hit compound to C0-C2 is exothermic (ΔH<0) and favorable reaction (ΔG<0) (see **Table 1**).

### TR-FRET: Functional characterization of compounds on actin-cMyBP-C binding

Compound-induced changes in TR-F and ITC results showing direct binding of the Hit compounds to C0-C2 suggested that the Hit compounds bind to C0-C2 and this modulates its binding with actin. We attempted to confirm effects on binding using a standard cosedimentation assay where F-actin was pelleted by centrifugation and the amount of C0-C2 bound to the F-actin in the pellet was quantitated. Unfortunately, all 3 Hit compounds caused C0-C2 to non-specifically (in the absence of any actin) bind to the centrifuge tubes used for the assay. This was the case even though the tubes are pre-coated with BSA to prevent non-specific sticking.

Therefore, we performed an intermolecular FRET assay to follow C0-C2 binding to actin (**Fig. 5A**). Actin was labeled with the fluorescent donor fluorescein-5-maleimide (FMAL) and incubated with increasing concentrations of C0-C2 labeled with the acceptor tetramethylrhodamine (TMR) at amino acid Cys225 (see Experimental Procedures) in the C1 domain. Binding of TMR-C0-C2 to FMAL-actin resulted in TR-FRET (**Fig. 5B** and **Fig. S1B** in *Supplemental Figures*). TR-FRET binding curves for nonphosphorylated and phosphorylated C0-C2 showed that this TR-FRET was dose-dependent (**Fig. 5C**). These binding curves with K_d_’s around 1 and 3 μM are in good agreement with the TR-F generated binding curves and those obtained by cosedimentation (7). At 1 μM FMAL-actin plus 2 μM TMR-C0-C2 (the same conditions as the original screen and for the dose response curves in **Fig. 3A**) 20 μM suramin and ATA and 50 μM NF023 almost completely eliminated TR-FRET measured binding of both nonphosphorylated and phosphorylated C0-C2 (**Fig. 5D**).

**Figure 5.**
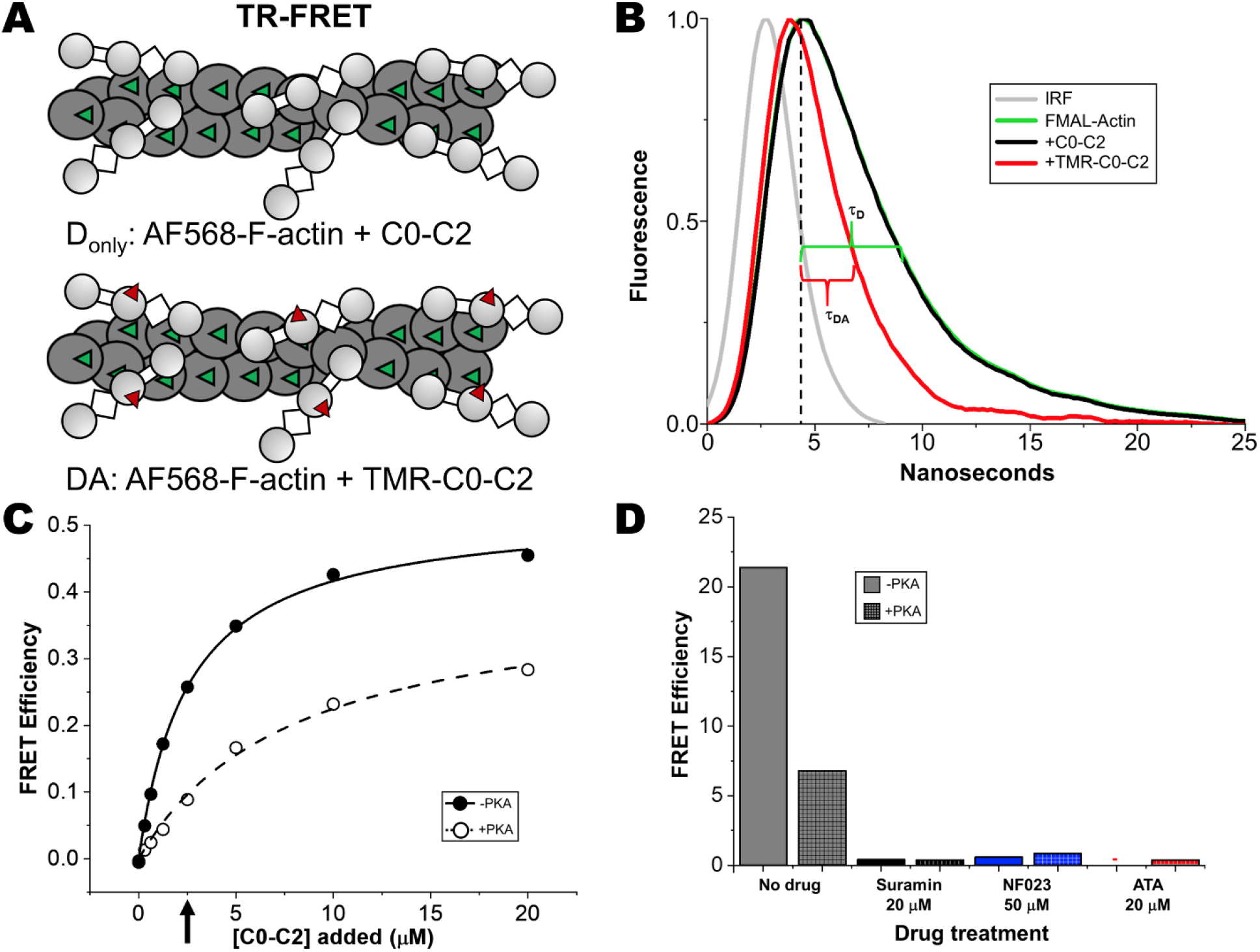
TR-FRET binding curve of PKA and Hit compounds. ***(A)*** Cartoon depiction of F-actin labeled with FMAL (green triangles) bound to C0-C2 (unlabeled), the donor-only sample (D_only_). Actual labeling is only 10%; 1 FMAL for every 10 actin monomers. And the same bound to C0-C2^Cys225^ labeled with TMR (red triangles), the donor-acceptor sample (DA). ***(B)*** Fluorescence waveforms of FMAL-actin (green line) and the same in the presence of 20 μM C0-C2^Cys225^ (black line) and 20 μM TMR-C0-C2^Cys225^ (red line), all normalized to maximal fluorescence (waveforms without normalization can be found in **Fig. S1B** in *Supplemental Figures*). Lifetime (τ) is the time at which the peak (vertical dashed line) fluorescence decays to ~0.37 (1/e). These times are indicated by the green bracket (τ_D_ = 4.2 ns for FMAL-actin with or without bound unlabeled C0-C2) and red bracket (τ_DA_ = 2.0 ns for FMAL-actin binding to TMR-C0-C2^Cys225^). The instrument response function (IRF) is also shown (grey line). ***(C)*** TR-FRET-based binding curve of 1 μM FMAL-actin and 0-20 μM TMR-C0-C2^Cys225^ ±PKA. The concentration of C0-C2 (2 μM) used in in the original screen and in testing of compound effects is indicated with an arrow. ***(D)*** Effect of adding compounds to 1 μM FMAL-actin-2 μM TMR-C0-C2^Cys225^ ±PKA (arrow in *C*). Compound concentrations are indicated. Aurintriboxcylic acid (ATA) eliminated actin-C0-C2 FRET to near zero (red -). Data are means ± SE; however, errors are too small to be seen outside of the data points or bar borders. n=9-10 for the binding curve and n=4-10 for the compound tests from 2 separate actin and C0-C2 preparations. All values are the average ± SE.

### TPA: Characterization of Hit compounds’ effects on actin structural dynamics

The 3 Hit compounds might interact directly with actin as well as with C0-C2. ITC showed only weak interaction of F-actin with suramin and no interaction with NF023 or ATA. We tested this further by monitoring compound effects on F-actin’s structural dynamics by using transient phosphorescence anisotropy (TPA) of actin labeled at Cys-374 with erythrosine iodoacetamide (ErIA) (**Fig. 6A**). None of the 3 compounds we identified in this screen affected actin’s structural dynamics as measured by TPA. For comparison we show a 4^th^ compound, GNF-5, that does bind actin and changes its anisotropy (**Fig. 6B**). This control was one of the compounds removed from the analysis due to effects on actin-alone in the screen.

**Figure 6.**
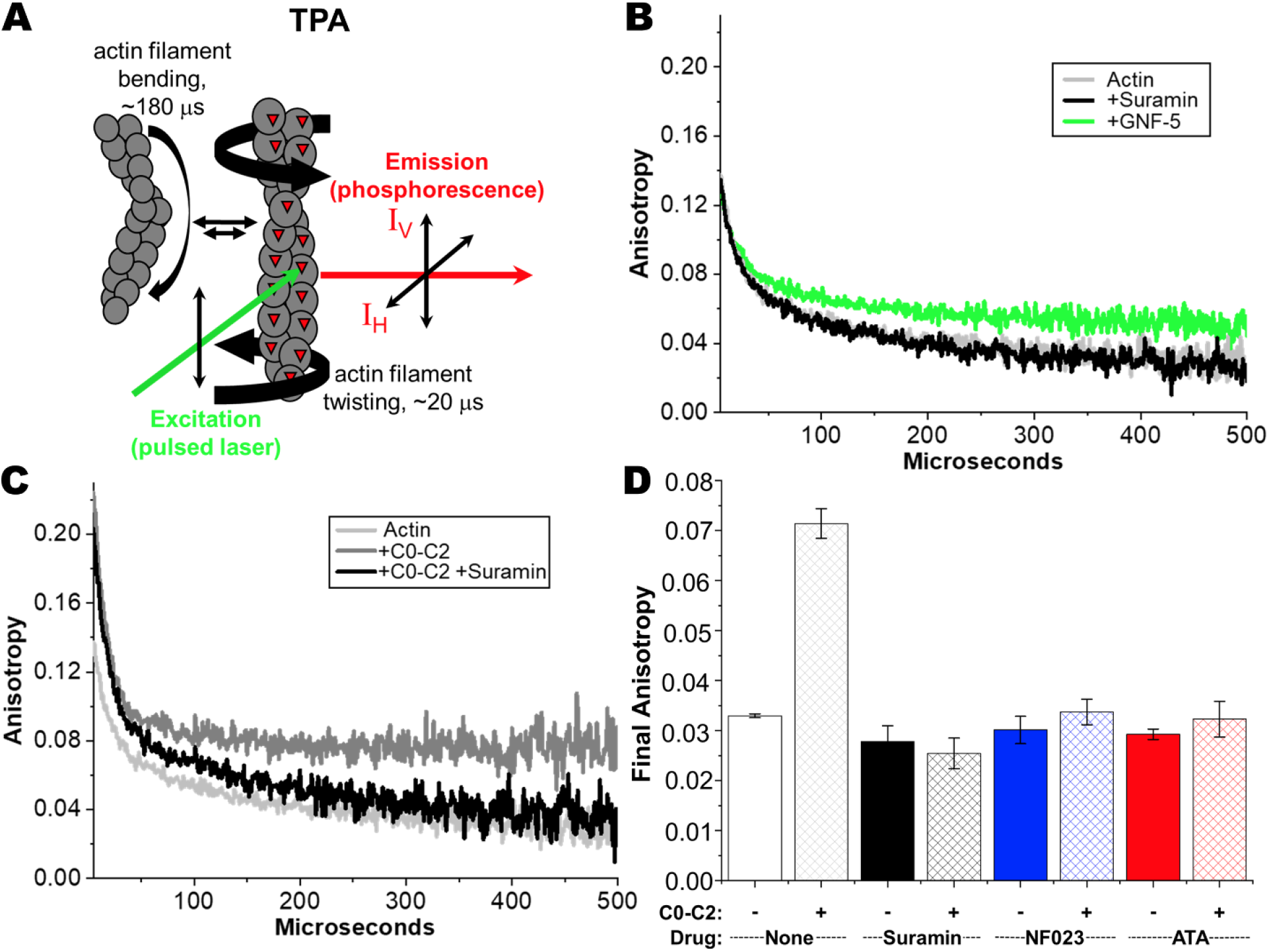
Effects of Hit compounds on actin rotational dynamics. ***(A)*** Schematic of transient phosphorescence anisotropy (TPA) measurement of actin labeled with erythrosine iodoacetamide (ErIA) where anisotropy of the emitted phosphorescence changes with the degree of actin filament twisting (from Bunch *et al*. 2019 *J Biol Chem*.) ***(B)*** TPA of ErIA-actin alone (light grey) +Suramin (black) or +GNF-5 (green). ***(C)*** TPA of actin alone (light grey) actin+C0-C2 (dark grey) actin+C0-C2+Suramin (black). ***(D)*** Effects of Hit compounds on TPA of actin (solid bars) or actin+C0-C2 (criss-cross bars). Compounds were present at 20 μM for Suramin and ATA, and 50 μM for NF023.

This TPA assay was used to test the 3 Hit compounds for their ability to inhibit C0-C2 binding to actin. We previously showed that actin structural dynamics are restricted by C0-C2 binding (10,11). **Fig. 6C** shows the effect of C0-C2 binding on F-actin’s final anisotropy (detecting rotational motions). Actin’s final anisotropy was increased from 0.033 ± 0.001 to 0.071 ± 0.003 in the presence of C0-C2. Addition of the 3 Hit compounds completely removes the C0-C2 effects on F-actin anisotropy (returning to ~0.033, **Fig. 6C-D**), consistent with the compounds binding to C0-C2 and preventing it from binding to actin.

## Discussion

The recently described TR-F assay, which monitors cMyBP-C’s amino terminal C0-C2 domains binding to actin (7), successfully identified the first three compounds capable of inhibiting C0-C2 interactions with actin. Characterization of this inhibition using a new TR-FRET assay, ITC, and TPA indicate that these 3 compounds bind directly to C0-C2, inhibiting its ability to bind to actin. Thus, we have identified the first 3 cMyBP-C-binding compounds and validated their ability to eliminate cMyBP-C’s actin-binding function in orthogonal assays.

Three compounds from ~1000 were reproducibly identified in 2 full screens of the LOPAC. That only 3 of the top 25 were reproducibly identified demonstrates the importance of replicate screening. 22 of the top preliminary hits in each screen did not show large effects in the other replicate screen and are considered false positives. In 2 additional screens of the same library, though not complete or optimal, the same 3 compounds but no others were found in the top 25 preliminary hits. We conclude that duplicate screening is necessary and sufficient for reliably identifying compounds using this screening technique. The reproducible Hit rate of ~0.3% suggests that screening of larger libraries of compounds will yield a manageable number of compounds for further characterization.

Further characterization was needed to determine whether the identified compounds inhibit C0-C2-actin interactions by binding to one or both proteins. All 3 compounds were found to bind C0-C2 and not actin by using a combination of ITC, TR-FRET, and TPA assays.

ITC showed clear binding to C0-C2 in the absence of actin but did not demonstrate binding to actin at the appropriate concentration. K_d_ values determined for Hit compound binding were similar to the half-maximal inhibition values determined by dose-response curves using the original TR-F assay. The two assays showed a consistent ranking of activity with suramin>ATA>NF023 for binding and inhibition. The half-maximal inhibition values were roughly 2-3 times lower than the K_d_ values. We have not investigated this further but note that the n value of the ITC results (**Table 1**) suggests that 2-3 molecules of the compounds bind to each C0-C2. If inhibition is achieved by the binding of only one molecule of compound, this could explain the difference between the ITC K_d_ and TR-F half-maximal inhibition values.

Adding a FRET acceptor (TMR) to C0-C2 allowed us to monitor its interaction with F-actin labeled with donor (FMAL) by TR-FRET. Using this new assay, we were able to observe the anticipated reduced binding when C0-C2 was phosphorylated with PKA. This assay confirmed the TR-F results, showing dramatic reduction of nonphosphorylated or phosphorylated C0-C2-actin interactions by all 3 compounds. Use of conventional cosedimentation assays to confirm binding was hindered by the compounds causing high background values, as they promoted C0-C2 binding to the tubes used in this assay.

In addition to ITC, we assayed interactions of the 3 compounds with actin using TPA. None of the 3 compounds altered actin’s anisotropy, suggesting that they were not binding to actin. C0-C2 binding to F-actin increased its anisotropy as expected and all 3 compounds prevented this. This further confirmed that the compound effects were due to their binding to C0-C2 and preventing its binding to actin. This result contrasts with our results screening for compounds that modulate actin-myosin interactions (12), where most of the identified compounds had a direct impact on the TPA of actin alone.

Compounds binding to cardiac MyBP-C but not to actin may be expected to have fewer side-effects than those binding to the very well conserved actin present in all cells. It will be important to determine whether these (and future) compounds that bind to cardiac MyBP-C also bind to skeletal MyBP-C. The observations from ITC that these compounds may bind to C0-C2 in a 2-3:1 ratio suggest that they bind to multiple MyBP-C domains such as C0, C1 and C2. Finally, it will be important to characterize the compounds’ effects on C0-C2 binding to myosin.

Much remains to be learned about the effects of these compounds on cardiac muscle contractility, but the approach used in this work clearly indicates that the TR-F assay is suitable for screening thousands of compounds to identify cMyBP-C binding Hits capable of altering its function. We now have, for the first time, a validated high-throughput screen focused on cMyBP-C, a known key factor in heart failure.

## Experimental Procedures

### Actin preparations and labeling

Actin was prepared from rabbit skeletal muscle by extracting acetone powder in cold water as described in Bunch *et al*. (13).

For time-resolved fluorescence experiments (TR-F), actin was labeled at Cys-374 with Alexa Fluor™ 568 C5 maleimide (AF568, ThermoFisher Scientific, Waltham, MA) and in preliminary experiments with Alexa Fluor™ 546 C5 maleimide (AF546). Labeling was done on F-actin. To 50 μM of G-actin in G-buffer (10 mM Tris pH 7.5, 0.2 mM CaCl_2_, 0.2 mM ATP), Tris pH 7.5 was added to a final concentration of 20 mM and actin was then polymerized by the addition of 3 M KCl (to a final concentration of 100 mM) and 0.5 M MgCl_2_ (to a final concentration of 2 mM), followed by incubation at 23 °C for 1 hour. AF568 was added to a final concentration of 50-100 μM (from a 20 mM stock in DMF). Labeling was done for 1 hour at 23 °C and then overnight at 4 °C. For TR-FRET experiments, actin was similarly labeled with fluorescein-5-maleimide (FMAL). Labeling with FMAL was done at a final FMAL concentration of 1 mM for 5 hours at 23 °C and then overnight at 4 °C. Labeling was stopped by the addition of a five-fold molar excess of DTT. Unincorporated dye (AF568 or FMAL) was removed by cycling the actin through F-actin and G-actin states as described in (7). For FMAL-labeled actin, to avoid TR-FRET between FMAL on neighboring Cys-374 residues of actin monomers in F-actin, unlabeled G-actin was mixed with FMAL-actin to achieve 10% FMAL-actin prior to the final actin polymerization. For phosphorescence experiments (TPA), actin was labeled at Cys374 with erythrosin-5’-iodoacetamide (ErIA; AnaSpec, Fremont, CA)(11,14). Fluorescent and phosphorescent labeled F-actins were stabilized by the addition of equimolar phalloidin.

For all assays F-actin was resuspended in and/or dialyzed against MOPS-actin binding buffer, M-ABB (100 mM KCl, 10 mM MOPS pH 6.8, 2 mM MgCl_2_, 0.2 mM CaCl_2_, 0.2 mM ATP, 1 mM DTT).

### Recombinant human cMyBP-C fragment preparations and labeling

pET45b vectors encoding *E. coli* optimized codons for the C0-C2 portion of human cMyBP-C with N-terminal 6x His tag and TEV protease cleavage site were obtained from GenScript (Piscataway, NJ). For TR-FRET binding assays, we mutated C0-C2 so that it contained a single cysteine at position 225, a surface-exposed residue in the C1 domain. To achieve this, the 5 endogenous cysteines in C0-C2 were mutated to amino acids found in other MyBP-C proteins at analogous positions and then His-225 was mutated to Cys. This single cysteine containing C0-C2^Cys225^ is described in detail in Kanassataga et al. (manuscript in preparation). Mutations were engineered in the human cMyBP-C C0-C2 using a Q5 Site-Directed Mutagenesis Kit (New England Bio Labs, Ipswich, MA). All sequences were confirmed by DNA sequencing (Eton Biosciences, San Diego, CA). Protein production in *E. coli* BL21DE3-competent cells (New England Bio Labs, Ipswich, MA) and purification of C0-C2 protein using His60 Ni Superflow resin was done as described (13). C0-C2 (with His-tag removed by TEV protease digestion) was further purified using size-exclusion chromatography to achieve >90% intact C0-C2 as described (10) and then concentrated, dialyzed to 50/50 buffer (50 mM NaCl and 50 mM Tris, pH 7.5) and stored at 4 °C.

For TR-FRET experiments, C0-C2^Cys225^ was labeled with tetramethylrhodamine (TMR) in 50/50 buffer. C0-C2^Cys225^ (50 μM) was first treated with the reducing agent TCEP (200 μM) for 30 minutes at 23 °C, and then TMR was added (from a 20 mM stock in DMF) to a final concentration of 200 μM. Labeling was done for 1 hour at 23 °C and terminated by the addition DTT (to 1 mM). Unincorporated dye was removed by extensive dialysis against M-ABB buffer. The degree of labeling was 0.95 dye/C0-C2 as measured by UV-vis absorbance.

### In vitro phosphorylation of cMyBP-C

C0-C2 was treated with 7.5 ng PKA/μg C0-C2 at 30 °C for 30 min, as described in (7,10).

### Fluorescence data acquisition

Fluorescence lifetime measurements were acquired using a high-precision fluorescence lifetime plate reader (FLTPR; Fluorescence Innovations, Inc., Minneapolis, MN) (13,15), provided by Photonic Pharma LLC (Minneapolis, MN). For TR-F experiments, AF568 (or AF546)-labeled F-actin (alone or mixed with C0-C2) was excited with a 532-nm microchip laser (Teem Photonics, Meylan, France) and emission was filtered with a 586/20-nm band pass filter (Semrock, Rochester, NY). For TR-FRET experiments, FMAL was excited with a 473-nm microchip laser (Bright Solutions, Cura Carpignano, Italy) and emission was filtered with 488-nm long pass and 517/20-nm bandpass filters (Semrock, Rochester, NY). The photomultiplier tube (PMT) voltage was adjusted so that the peak signals of the instrument response function (IRF) and the TR-F biosensor were similar. The observed TR-F waveform was analyzed as described previously (7,16).

### LOPAC library screen

The 1280 LOPAC compounds were received in 96-well plates and were reformatted into 1,536-well flat, black-bottom polypropylene plates (Greiner Bio-One, Kremsmunster, Austria). 50 nl of compounds were dispensed using an automated Echo 550 acoustic liquid dispenser (Labcyte, Sunnyvale, CA). LOPAC compounds were formatted into plates, with the first two and last two columns loaded with 50 nl of DMSO and used for compound-free controls. The final concentration of the compounds was 10 μM. These assay plates were then heat-sealed using a PlateLoc Thermal Microplate Sealer (Agilent Technologies, Santa Clara, CA) and stored at −20 °C. Before screening, compound plates were equilibrated to room temperature (25 °C). 1 μM AF568-labeled actin without or with 2 μM C0-C2 in M-ABB was dispensed by a Multidrop Combi Reagent Dispenser (ThermoFisher Scientific, Waltham, MA) into the 1,536-well assay plates containing the compounds. Plates were incubated at room temperature for 20 min before recording the data with the FLTPR. Plates were re-scanned after 120 min incubation.

Two full screens were done using different batches of 568-labeled actin, C0-C2, and test compounds. Two additional sub-optimal (due to liquid dispensing errors that necessitated removing 191 samples from the analysis, and a batch of 568-actin that showed lower than expected lifetime changes with the binding of C0-C2 in the absence of compounds) screens were included in the analysis (**Tables S3** and **S4** in *Supplemental Tables*) as they yielded complementary results.

### TR-F and HTS data analysis

Following data acquisition, TR-F waveforms observed for each well were convolved with the IRF to determine the lifetime (τ) (Eq. 1) by fitting to a single-exponential decay (7,16).

The decay of the excited state of the fluorescent dye attached to actin at Cys-374 to the ground state is:

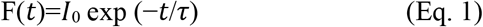

where *I_0_* is the fluorescence intensity upon peak excitation (*t* = 0) and τ is the fluorescence lifetime (t = τ when *I* decays to 1/e or ~37% of *I_0_*).

TR-F assay quality was determined from controls (DMSO-only samples) on each plate as indexed by the *Z’* factor. A value of 0.5 to 1 indicates excellent assay quality while 0.0-0.5 indicates a doable assay quality (7,17,18):

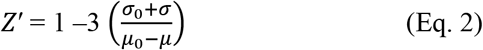

where σ_0_ and σ are the standard deviations (SD) of the controls τ_0_ and τ, respectively, and μ_0_ and μ are the means of the controls τ_0_ and τ, respectively. Here, the two-sample comparison was AF568-actin with DMSO (τ_0_) versus AF568-actin with C0-C2 and DMSO (τ).

A compound was considered a Hit if it was in the top 25 compounds in each of two independent screenings of the LOPAC. Top compounds were those that increased the lifetime of AF568-actin when C0-C2 was present (indicating inhibition of C0-C2 binding). Compounds that altered AF568-actin lifetime when C0-C2 was not present by >4 S.D. relative to that of the control samples (those exposed to 0.1% DMSO) were first removed from the analysis. This resulted in the removal, from analysis, of 210 compounds from the first screen and 106 compounds from the second screen. Any change in lifetime induced by a compound on AF568-actin alone was then subtracted from the effect of the compound on AF568-actin when C0-C2 was present. The resulting C0-C2-dependent change for all compounds was ranked.

### TR-F concentration-response assay

The Hit compounds were dissolved in DMSO to make a 10 mM stock solution, which was serially diluted in in 96-well mother plates. Hits were screened at eight concentrations (0.5 to 100 μM). Compounds (1 μl) were transferred from the mother plates into 384-well plates using a Mosquito HV liquid handler (TTP Labtech Ltd., Hertfordshire, UK). The same procedure of dispensing as for the pilot screening was applied in the TR-F concentration-response assays. Concentration dependence of the TR-F change was fitted using the Hill equation (19):

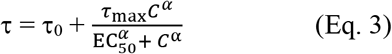

where τ and τ_0_ are TR-F in the presence and in the absence of the compound, τ_max_ is the maximum effect, *C* is the compound concentration, EC_50_ is the compound concentration for which 50% of maximum effect is obtained, and α is the Hill coefficient of sigmoidicity.

### Isothermal Titration Calorimetry (ITC)

The titration of compounds binding to cMyBP-C was performed on an Affinity ITC (TA Instruments, New Castle, DE). Each compound in M-ABB buffer (2,000-3,000 μM) was titrated into C0-C2 in M-ABB buffer (100 μM, 350 μl). For the titration, a preliminary 2.5-μl injection (ignored in data analysis) was followed by 19-39 subsequent 2.5-μl injections with 4 min waits between each injection. The reference cell was filled with distilled water. To account for heat changes of dilution and mechanical mixing, binding data were corrected by subtraction of the final plateau injection values (when binding is complete) prior to curve fitting. Corrected binding data was processed using the NanoAnalyze analysis software (TA Instruments, New Castle, DE).

### FMAL-Actin-TMR-C0-C2 TR-FRET binding assay

1μM F-Actin labeled (to 10%) on Cys-374 with FMAL (donor fluor) was incubated with increasing concentrations (0-20 μM) of C0-C2^Cys225^ unlabeled (for donor only (D) data acquisition) or labeled to 95% with TMR (acceptor fluor) (for donor-acceptor (DA) data). Lifetimes of FMAL-actin were determined for D and DA at each concentration of C0-C2^Cys225^. TR-FRET efficiency (1 – (τ_DA_/τ_D_)) was determined for each C0-C2^Cys225^ concentration. The binding curve was generated from data from 2 separate preparations of actin and C0-C2^Cys225^. Testing of compound effects by TR-FRET was similarly done at 1 μM FMAL-actin with unlabeled C0-C2^Cys225^ (D) and TMR-C0-C2^Cys225^ (DA), both at 2 μM.

### Transient phosphorescence Anisotropy (TPA)

Phalloidin-stabilized ErIA-actin was prepared and diluted in M-ABB to 1 μM alone or in combination with each compound at concentrations that first exhibited maximal effect in the initial TR-F assay; 20 μM for suramin and ATA and 50 μM for NF023. The same mixtures were tested in the presence of 2 μM C0-C2. These were added together with glucose catalase/oxidase to prevent photobleaching, and TPA was performed at 25 °C to determine anisotropy values (10).

### Statistics

Average data are provided as mean ± standard error (SE) and each experiment was done with 2 separate protein preparations.

## Data Availability

All data discussed are presented within the article.

## Author Contributions

B.A.C., T.A.B., D.D.T., and P.G.T. designed the study and wrote the paper. T.A.B., V.C.L., and R.-S.K. purified recombinant proteins and DNA. T.A.B., V.C.L., P.G.T., and A.W. undertook purification of actin filaments for cosedimentation assays, TPA, TR-F, TR-FRET, and ITC assays. V.C.L. and T.A.B. conducted cosedimentation assays and T.A.B. and P.G.T. conducted TR-F assays. T.A.B. performed TR-FRET assays and ITC. R.-S.K. made the C0-C2^Cys225^ mutant and optimized its labeling. A.W. prepared compound plates. T.A.B. and B.A.C. analyzed cosedimentation assays, TPA, TR-FRET and ITC. T.A.B., B.A.C., P.G.T., A.R.T., and D.D.T. analyzed fluorescence lifetime data and assisted with preparation of the figures and tables. V.C.L. and A.R.T. assisted with critical evaluation of the results and edited the manuscript. All authors critically evaluated and approved the final version of the manuscript.

## FOOTNOTES

## Acknowledgements

This work was supported by NIH grants R37 AG26160 (to D.D.T.) and R01 HL141564 (to B.A.C.), and a University of Arizona Sarver Heart Center “Novel Research Project Award in the Area of Cardiovascular Disease and Medicine” (to B.A.C). We thank Samantha Yuen for technical assistance with the fluorescence lifetime plate reader.

## Abbreviations

DCM: dilated cardiomyopathy
HCM: hypertrophic cardiomyopathy
cMyBP-C: cardiac myosin-binding protein C
P/A: proline/alanine-rich linker between domains C0 and C1
M: M-domain, phosphorylatable linker between C1 and C2
C0-C2: N-terminal fragment of cMyBP-C comprised of C0-P/A-C1-M-C2 domains and linkers
HTS: high-throughput screen
FLTPR: fluorescence lifetime plate reader
DO: donor only
DA: donor plus acceptor
LOPAC: library of pharmacologically active compounds
AF568: Alexa Fluor 568
AF546: Alexa Fluor 546
F-actin: filamentous actin
G-actin: globular actin
TR-F: time-resolved fluorescence
TR-FRET: time-resolved fluorescence energy transfer
ITC: isothermal titration calorimetry
TPA: transient phosphorescence anisotropy
K_d_: dissociation constant
B_max_: maximum molar binding ratio
ATA: aurintricarboxylic acid
M-ABB: MOPS-actin binding buffer
DMSO: dimethylsulphoxide
DMF: dimethylformamide
BSA: bovine serum albumin
DTT: dithiothreitol
TCEP: tris(2-carboxyethyl)phosphine
ErIA: erythrosine iodoacetamide phosphorescent dye
PKA: protein kinase A
FMAL: fluorescein-5-maleimide
IAEDANS: 5-((((2-iodoacetyl)amino)ethyl)amino)naphtalene-1-sulfonic acid
TMR: tetramethylrhodamine
SD: standard deviations
SE: standard errors
DWR: direct waveform recording

