## Supplemental Materials for "High-throughput screen using fluorescence lifetime detects compounds that modulate myosin-binding protein C interactions with actin"

**Supplemental Material**

**Supplemental Tables**

**Table S1. Top 25 hits from Screen 1**

| | Rank | Row | Column | Actin-MyBP-C Drug Δτ (ns) | Actin Drug Δτ (ns) | Difference (ns) |
|---|---|---|---|---|---|---|
| **Suramin** | **1** | AD | 29 | 0.088 | -0.03 | 0.12 |
| **NF 023** | **2** | R | 21 | 0.058 | -0.03 | 0.09 |
| | **3** | R | 42 | 0.018 | -0.04 | 0.06 |
| | **4** | T | 27 | 0.008 | -0.04 | 0.05 |
| | **5** | R | 7 | 0.013 | -0.04 | 0.05 |
| | **6** | Q | 6 | 0.008 | -0.04 | 0.04 |
| **ATA** | **7** | K | 9 | 0.008 | -0.04 | 0.04 |
| | **8** | X | 2 | -0.003 | -0.04 | 0.04 |
| | **9** | G | 18 | -0.006 | -0.04 | 0.04 |
| | **10** | D | 34 | 0.011 | -0.02 | 0.03 |
| | **11** | Z | 2 | -0.010 | -0.04 | 0.03 |
| | **12** | R | 22 | -0.007 | -0.04 | 0.03 |
| | **13** | V | 25 | -0.008 | -0.04 | 0.03 |
| | **14** | D | 40 | 0.017 | -0.01 | 0.03 |
| | **15** | R | 30 | -0.006 | -0.04 | 0.03 |
| | **16** | V | 21 | -0.006 | -0.03 | 0.03 |
| | **17** | S | 37 | -0.005 | -0.03 | 0.03 |
| | **18** | T | 15 | -0.010 | -0.04 | 0.03 |
| | **19** | X | 7 | -0.005 | -0.03 | 0.03 |
| | **20** | K | 1 | 0.001 | -0.03 | 0.03 |
| | **21** | G | 40 | -0.005 | -0.03 | 0.03 |
| | **22** | X | 39 | -0.003 | -0.03 | 0.03 |
| | **23** | J | 21 | -0.014 | -0.04 | 0.03 |
| | **24** | R | 46 | -0.019 | -0.05 | 0.03 |
| | **25** | K | 19 | -0.017 | -0.04 | 0.02 |

Top 25 hits from Screen 1 ranked include 3 Hits: Suramin (grey), NF 023 (blue), and ATA (red). Compounds are ranked for largest Difference (ns) due to drug effects on the change in lifetime (Δτ, ns) for actin +C0-C2 after subtracting the effect of compounds on Δτ for actin alone. Location of compound in row and column of wells is listed.

**Table S2. Top 25 hits from Screen 2**

| | Rank | Row | Column | Actin-MyBP-C Drug Δτ (ns) | Actin Drug Δτ (ns) | Difference (ns) |
|---|---|---|---|---|---|---|
| **Suramin** | 1 | AD | 29 | 0.15 | 0.00 | 0.16 |
| | 2 | S | 5 | 0.11 | 0.00 | 0.11 |
| | 3 | T | 30 | 0.10 | 0.00 | 0.10 |
| | 4 | V | 38 | 0.09 | 0.00 | 0.10 |
| | 5 | S | 41 | 0.09 | 0.00 | 0.09 |
| | 6 | R | 28 | 0.08 | 0.00 | 0.08 |
| | 7 | T | 4 | 0.04 | -0.02 | 0.07 |
| | 8 | R | 12 | 0.05 | -0.01 | 0.07 |
| **NF 023** | 9 | R | 21 | 0.07 | 0.01 | 0.06 |
| | 10 | T | 20 | 0.04 | -0.02 | 0.06 |
| **ATA** | 11 | K | 9 | 0.06 | 0.00 | 0.06 |
| | 12 | R | 6 | 0.06 | 0.01 | 0.06 |
| | 13 | R | 4 | 0.03 | -0.03 | 0.06 |
| | 14 | T | 37 | 0.05 | 0.00 | 0.06 |
| | 15 | P | 40 | 0.06 | 0.01 | 0.05 |
| | 16 | T | 17 | 0.05 | 0.00 | 0.05 |
| | 17 | S | 16 | 0.04 | -0.01 | 0.05 |
| | 18 | T | 38 | 0.04 | -0.01 | 0.05 |
| | 19 | T | 18 | 0.04 | -0.01 | 0.05 |
| | 20 | T | 45 | 0.04 | -0.01 | 0.05 |
| | 21 | T | 13 | 0.05 | 0.00 | 0.05 |
| | 22 | R | 32 | 0.03 | -0.02 | 0.05 |
| | 23 | T | 19 | 0.03 | -0.01 | 0.04 |
| | 24 | W | 44 | 0.04 | -0.01 | 0.04 |
| | 25 | S | 20 | 0.02 | -0.02 | 0.04 |

Top 25 hits from Screen 2 ranked include 3 Hits: Suramin (grey), NF 023 (blue), and ATA (red). Compounds are ranked for largest Difference (ns) due to drug effects on the change in lifetime (Δτ, ns) for actin +C0-C2 after subtracting the effect of compounds on Δτ for actin alone. Location of compound in row and column of wells is listed.

**Table S3. Top 25 hits from Sub-optimal Screen 3**

| | Rank | Row | Column | Actin-MyBP-C Drug Δτ (ns) | Actin Drug Δτ (ns) | Difference (ns) |
|---|---|---|---|---|---|---|
| **Suramin** | **1** | AD | 29 | 0.18 | -0.03 | 0.21 |
| | **2** | M | 35 | 0.17 | -0.03 | 0.20 |
| **ATA** | **3** | K | 9 | 0.07 | -0.03 | 0.10 |
| | **4** | J | 18 | 0.08 | -0.02 | 0.10 |
| | **5** | M | 43 | 0.06 | -0.03 | 0.09 |
| | **6** | N | 40 | 0.07 | -0.03 | 0.09 |
| | **7** | N | 46 | 0.04 | -0.05 | 0.09 |
| | **8** | J | 12 | 0.06 | -0.02 | 0.09 |
| **NF 023** | **9** | R | 21 | 0.06 | -0.02 | 0.08 |
| | **10** | J | 10 | 0.05 | -0.03 | 0.08 |
| | **11** | AA | 47 | 0.03 | -0.04 | 0.08 |
| | **12** | AF | 13 | 0.03 | -0.03 | 0.07 |
| | **13** | M | 36 | 0.05 | -0.01 | 0.07 |
| | **14** | AA | 18 | 0.02 | -0.05 | 0.07 |
| | **15** | K | 45 | 0.05 | -0.02 | 0.06 |
| | **16** | J | 13 | 0.03 | -0.03 | 0.06 |
| | **17** | L | 10 | 0.05 | -0.01 | 0.06 |
| | **18** | N | 30 | 0.04 | -0.03 | 0.06 |
| | **19** | N | 34 | 0.02 | -0.04 | 0.06 |
| | **20** | AB | 47 | 0.02 | -0.04 | 0.06 |
| | **21** | AB | 12 | 0.04 | -0.02 | 0.06 |
| | **22** | AB | 29 | 0.02 | -0.04 | 0.06 |
| | **23** | N | 20 | 0.03 | -0.03 | 0.06 |
| | **24** | AF | 19 | 0.02 | -0.04 | 0.06 |
| | **25** | J | 38 | 0.03 | -0.03 | 0.06 |

Top 25 hits from Sub-optimal Screen 3 ranked include 3 Hits: Suramin (grey), NF 023 (blue), and ATA (red). Compounds are ranked for largest Difference (ns) due to drug effects on the change in lifetime (Δτ, ns) for actin +C0-C2 after subtracting the effect of compounds on Δτ for actin alone. Location of compound in row and column of wells is listed.

**Table S4. Top 25 hits from Sub-optimal Screen 4**

| | Rank | Row | Column | Actin-MyBP-C Drug Δτ (ns) | Actin Drug Δτ (ns) | Difference (ns) |
|---|---|---|---|---|---|---|
| **Suramin** | **1** | AD | 29 | 0.13 | 0.04 | 0.09 |
| **ATA** | **2** | K | 9 | 0.07 | 0.00 | 0.07 |
| | **3** | A | 3 | 0.07 | 0.01 | 0.06 |
| | **4** | A | 1 | 0.06 | 0.01 | 0.05 |
| | **5** | B | 12 | 0.01 | -0.03 | 0.04 |
| | **6** | B | 11 | 0.01 | -0.03 | 0.04 |
| | **7** | I | 11 | 0.02 | -0.02 | 0.04 |
| | **8** | I | 17 | 0.04 | 0.00 | 0.04 |
| | **9** | J | 21 | 0.03 | 0.00 | 0.04 |
| | **10** | M | 21 | 0.05 | 0.01 | 0.04 |
| | **11** | L | 15 | 0.05 | 0.01 | 0.04 |
| | **12** | P | 22 | 0.07 | 0.03 | 0.04 |
| | **13** | A | 4 | 0.03 | 0.00 | 0.04 |
| | **14** | B | 26 | 0.02 | -0.01 | 0.03 |
| | **15** | P | 43 | 0.02 | -0.01 | 0.03 |
| | **16** | L | 46 | 0.01 | -0.02 | 0.03 |
| | **17** | O | 42 | 0.01 | -0.02 | 0.03 |
| | **18** | L | 39 | 0.04 | 0.01 | 0.03 |
| | **19** | O | 5 | 0.06 | 0.03 | 0.03 |
| | **20** | B | 14 | 0.02 | -0.01 | 0.03 |
| **NF 023** | **21** | R | 21 | 0.04 | 0.01 | 0.03 |
| | **22** | P | 28 | 0.03 | 0.00 | 0.03 |
| | **23** | L | 33 | 0.04 | 0.01 | 0.03 |
| | **24** | M | 19 | 0.04 | 0.01 | 0.03 |
| | **25** | N | 30 | 0.05 | 0.02 | 0.03 |

Top 25 hits from Sub-Optimal Screen 4 ranked include 3 Hits: Suramin (grey), NF 023 (blue), and ATA (red). Compounds are ranked for largest Difference (ns) due to drug effects on the change in lifetime (Δτ, ns) for actin +C0-C2 after subtracting the effect of compounds on Δτ for actin alone. Location of compound in row and column of wells is listed.

**Supplemental Figures**

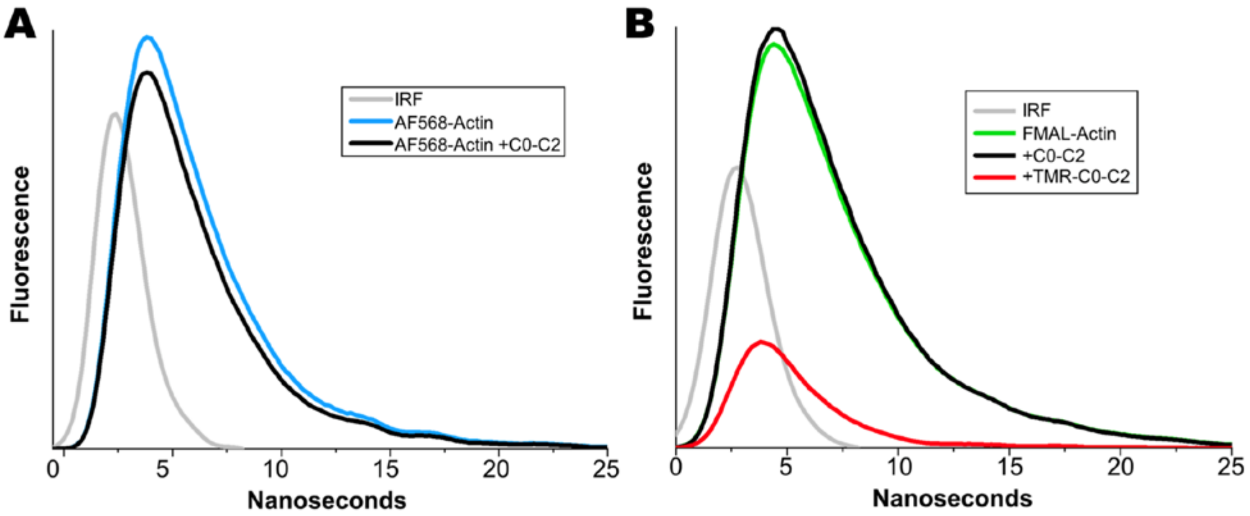


**Figure S1. TR-F and TR-FRET waveforms as detected by direct waveform recording (DWR).** ***(A)*** Waveforms of AF568-Actin ± C0-C2 (without normalization of fluorescence) detected in TR-F plate reader experiments. Normalized TR-F waveforms are found in **Fig. 2B** of main text. ***(B)*** Waveforms of donor-only, FMAL-Actin ± unlabeled C0-C2, and donor-acceptor, FMAL-Actin +TMR-C0-C2 (without normalization of fluorescence) detected in TR-FRET plate reader experiments. FRET occurs in donor-acceptor waveform (red) where both intensity and lifetime are reduced due to acceptor quenching of donor. Normalized TR-FRET waveforms are found in **Fig. 5B**. The instrument response function (IRF) is shown in grey.

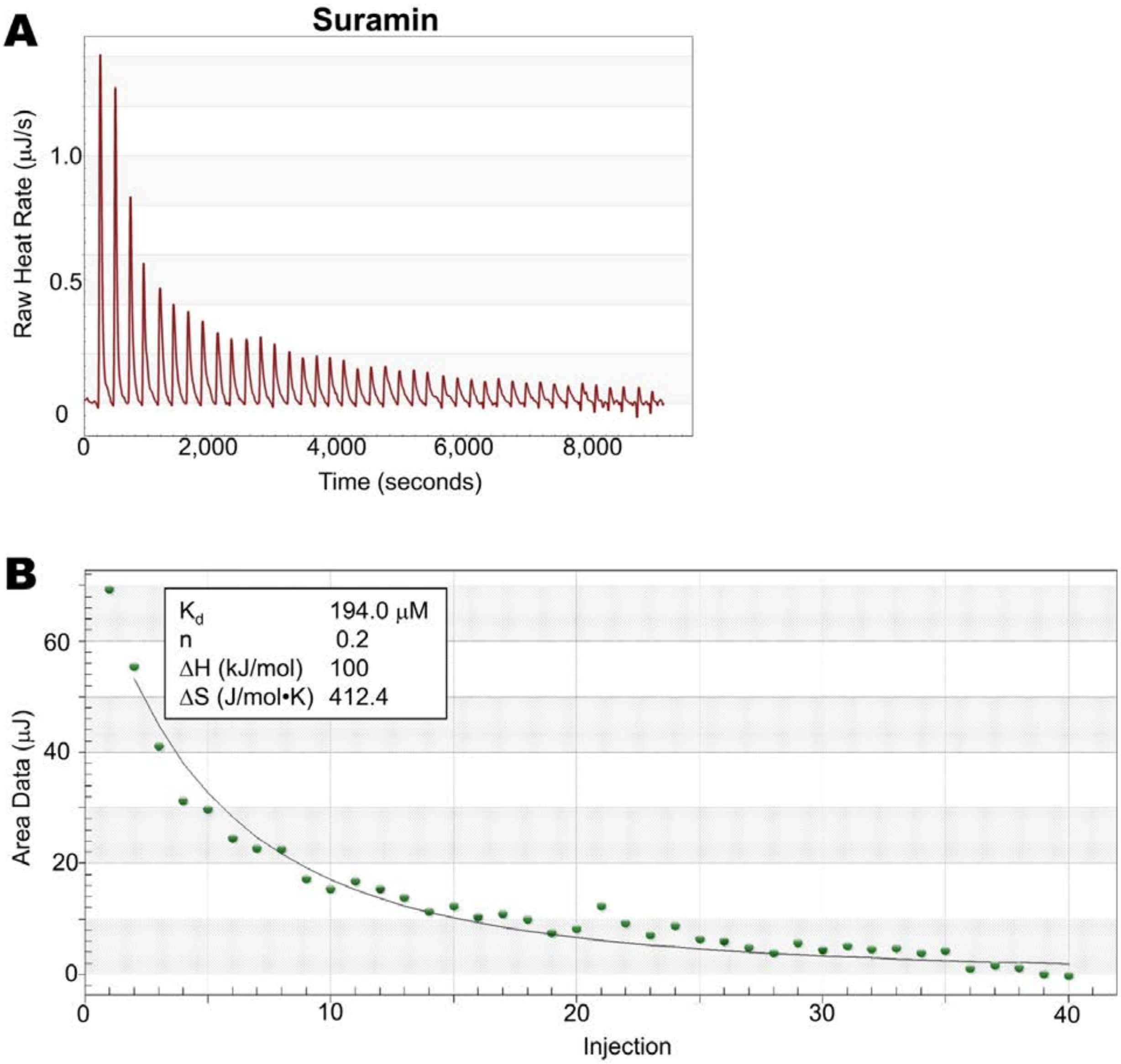


**Figure S2**. ITC of Suramin binding to actin alone. ***(A)*** Titration of 100 μM F-actin (in 350 μl) with 40 2.5 μl injections of 3,000 μM Suramin. ***(B)*** Fitted ITC binding curve for Suramin binding. Binding to actin was not apparent for NF023 or ATA ($-0.2 <$ Raw Heat Rate (μJ/s) $< 0.2$), but weak interactions were observed for Suramin with actin alone ($K_d$ ~200 μM).
